# Selective participation of single cortical neurons in neuronal avalanches

**DOI:** 10.1101/2020.10.21.349340

**Authors:** Timothy Bellay, Woodrow L. Shew, Shan Yu, Jessica J. Falco-Walter, Dietmar Plenz

**Affiliations:** Section on Critical Brain Dynamics, National Institute of Mental Health, National Institutes of Health, Bethesda, MD, USA; Department of Neuroscience, Brown University, Providence, RI, USA; Department of Physics, University of Arkansas, AR, USA; Brainnetome Center, Inst. of Automation, Chinese Academy of Sciences, China; Department of Neurology, Stanford University, Palo Alto, CA, USA

**Author notes:** These authors contributed equally to this work (Timothy Bellay, Woodrow L. Shew). Correspondence: Dietmar Plenz, Ph.D., Section on Critical Brain Dynamics, National Institute of, Mental Health, Porter Neuroscience Research Center, Rm 3A-1000, 35 Convent Drive, Bethesda, MD 20892.

**Keywords:** Nonhuman primate, rat, prefrontal cortex, primary motor cortex, high-density microelectrode array, local field potential, whole-cell patch recording, resting activity, cell assemblies

## Abstract

Neuronal avalanches are scale-invariant neuronal population activity patterns in cortex that emerge *in vivo* in the awake state and *in vitro* during balanced excitation and inhibition. Theory and experiments suggest that avalanches indicate a state of cortex that improves numerous aspects of information processing by allowing for the transient and selective formation of local as well as system-wide spanning neuronal groups. If avalanches are indeed involved with information processing, one might expect that particular single neurons would participate in particular avalanche patterns selectively. Alternatively, all neurons could participate with equal likelihood in each avalanche as would be expected for a population rate code. Distinguishing these hypotheses, however, has been difficult as robust avalanche analysis requires technically challenging measures of their intricate organization in space and time at the population level, while also recording sub- or suprathreshold activity from individual neurons with high temporal resolution. Here we identify repeated avalanches in the ongoing local field potential (LFP) measured with high-density microelectrode arrays in the cortex of awake nonhuman primates and in acute cortex slices from rats. We studied extracellular unit firing *in vivo* and intracellular responses of pyramidal neurons *in vitro*. We found that single neurons participate selectively in specific LFP-based avalanche patterns. Furthermore, we show *in vitro* that manipulating the balance of excitation and inhibition abolishes this selectivity. Our results support the view that avalanches represent the selective, scale-invariant formation of neuronal groups in line with the idea of Hebbian cell assemblies underlying cortical information processing.

## Introduction

Understanding how the collective dynamics of large cortical networks emerge from the activities of single neurons is one of the fundamental challenges in systems neuroscience. Given the limitations of accurately recording from most neurons in real cortical networks, this challenge has been typically reduced to how the collective dynamics *measured* in large neuronal networks relate to the activity of single neurons. Of particular interest in this context has been the discovery of ‘neuronal avalanches’ in spontaneous (Beggs and Plenz, 2003;Petermann et al., 2009;Miller et al., 2019) and evoked cortical activity (Shew et al., 2015;Yu et al., 2017) in which the local collective dynamics of cortex has been mapped using the local field potential (LFP). More specifically, it has been reliably found for slice cultures, acute slices, rodents and nonhuman primates that the spatial and temporal spread of transient and fast deflections in the cortical LFP (Yu et al., 2014), when tracked using high-density microelectrode arrays (MEAs), obeys a power law relationships in the size of LFP patterns, the hallmark of avalanches.

The power law in avalanche size demonstrates that large LFP patterns, i.e. those that engage most of the area monitored by the electrode array, are significantly more common than expected by chance (Yu et al., 2014). Based on hierarchical clustering, the diverse organization of LFP avalanches *in vitro* was found to organize into families of similar spatial patterns, i.e. avalanche families (Beggs and Plenz, 2004;Stewart and Plenz, 2006). The commonality in large spatial patterns organized into families then raises the question whether individual neurons participate in the generation of avalanches or avalanche families selectively, or, alternatively, in a random manner. In case of the former, each single neuron’s response would correlate preferential with some avalanche families, whereas in the latter case, all single neuron responses would correlate equally with avalanches at a rate proportional to avalanche size. Recent cellular analysis using 2-photon imaging has demonstrated that the spontaneous and evoked firing in groups of cortical neurons in the awake rodent organizes as scale-invariant avalanches (Bellay et al., 2015;Karimipanah et al., 2017;Bowen et al., 2019;Ribeiro et al., 2020). Similarly, extracellular unit recordings in the rodent during wakefulness, exploration, and sleep identified state specific and repeated spike avalanche patterns (Ribeiro et al., 2016). Yet, it is currently not known how supra- and subthreshold activity of individual neurons relate to the diversity of LFP avalanches.

Here we studied the relationship between avalanches and single-neuron activity by comparing multi-site LFP recordings with simultaneously measured extra- and intracellular activity of single neurons. More specifically, when a spatial pattern of the LFP was found to repeat during a recording, we searched for reliable recruitment of single neurons during each repeated occurrence. First, we studied extracellular unit activity and LFP signals recorded during ongoing activity from layers 2/3 of premotor cortex in awake monkeys. Since it is not feasible to separate the effects of local and distant sources of the LFP in awake animals, we next carried out complementary studies in acute slices of rat cortex, for which the origins of the LFP signals are intrinsic to the cortex. For the slice studies, we combined intracellular whole-cell patch recordings of single layer 2/3 pyramidal neurons with multi-site LFP recordings. In line with our hypothesis, both *in vivo* and *in vitro*, we found that neurons participate selectively and reliably in particular avalanche patterns. We further demonstrate that this selective relationship between neurons and avalanches requires intact synaptic inhibition.

## Material and Methods

### Nonhuman Primate Recordings

Two adult nonhuman primates (*Macaca mulatta*), 1 female (monkey 1; Victoria) and 1 male (monkey 2; Noma) were studied. High-density MEAs (96 electrodes, 10 × 10 grid configuration with no corner electrodes, 0.4-mm inter-electrode spacing and 1.0-mm electrode length; from BlackRock Microsystems, Salt Lake City, UT) were chronically implanted in the arm representation region of the left pre-motor cortex. Recordings were done at least one week following surgery. Ongoing activity was recorded for a period of 30 min, during which the monkey was awake but did not perform any behavioural task. Extracellular signals were recorded at 30 kHz. In post-recording processing, LFP signals were downsampled to 500 Hz and band-pass filtered at 1 – 100 Hz. One exception was the analysis presented in Figure 1D, for which the band-pass was 3 – 100 Hz.

**Figure 1.**
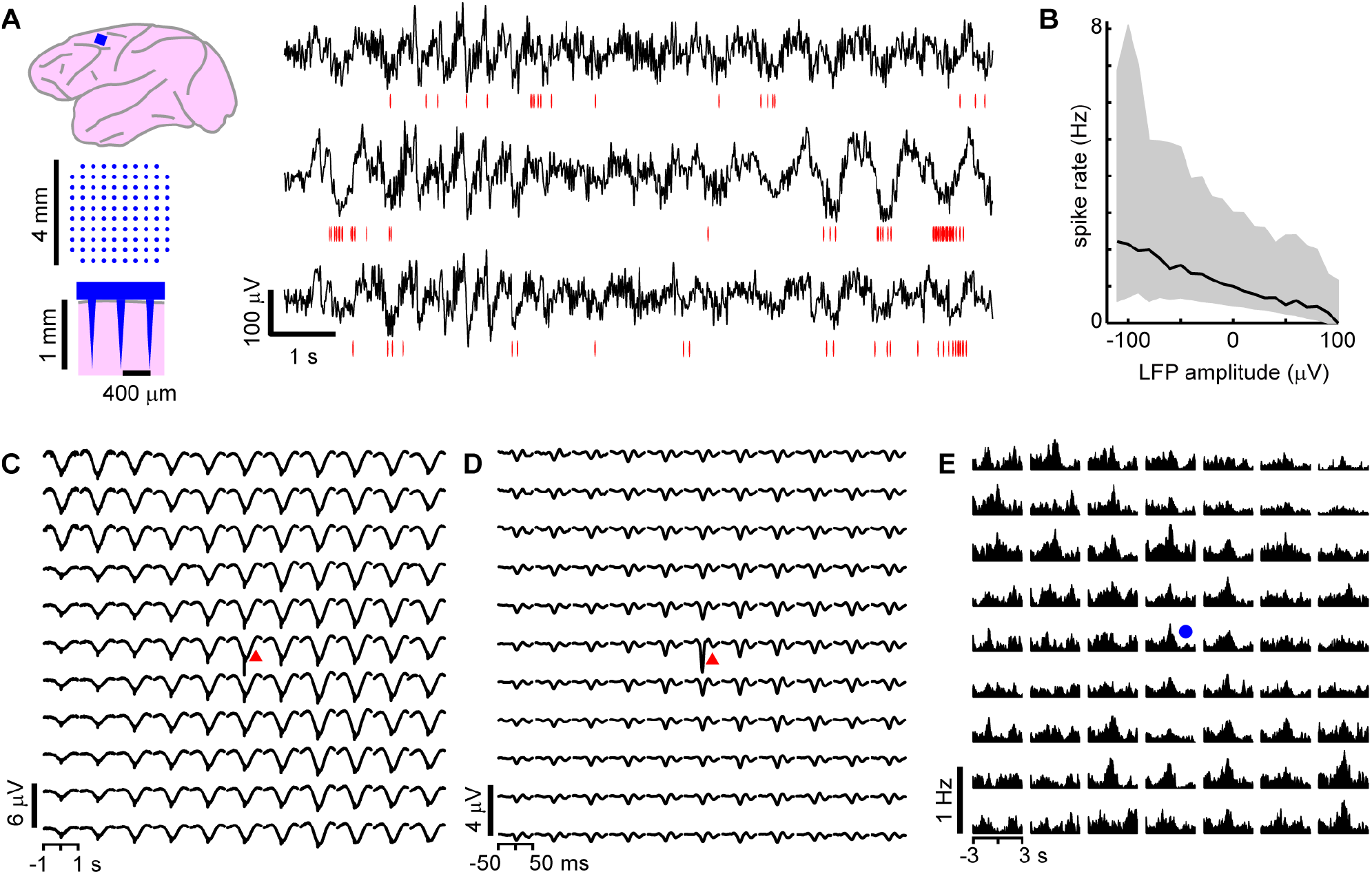
On average, local unit activity associates with negative LFP deflections over spatially extended cortical area in monkey cortex. **A.** *Left*: A 96 channel electrode array (blue) was implanted within superficial layers of premotor cortex in two macaque monkeys. *Right*: Simultaneous recordings of awake-state ongoing LFP (black) and single unit activity (red – spike times) from three recording sites. Units exhibit increased firing around the time of negative LFP deflections. **B.** Unit activity increases with negative LFP amplitude. Unit spike rate as a function of LFP amplitude computed for all times (in consecutive 50 ms windows). Units and LFP recorded at the same site were compared. Displayed is the average over all sites and units for both monkeys. Large deviations from the average relationship are typical (shaded region – lower to upper quartile, line – median). **C.** Spike-triggered average LFP waveforms indicate that widespread, slow negative LFP fluctuations are associated with local spiking (red triangle indicates site of local unit). LFP was band-pass filtered between 1 – 100 Hz. **D.** Same as in C, but with low frequency components of the LFP removed (band-pass, 3 – 100 Hz), spike-triggered average LFP waveforms indicate that local, sharp negative LFP deflections are typically associated with spiking as reported previously (Petermann et al., 2009). **E.** The peak times of large amplitude negative LFP deflections (nLFPs; bandpass filter, 1 – 100 Hz) were used to compute nLFP-triggered average spike count histograms. Consistent with C, units both near to and distant from the trigger site (blue dot) exhibited significant increases in firing with no clear decaying relationship over distance. For C, D and E all units were compared to all LFP recordings and then averaged together keeping track of relative locations of the unit with respect to the LFP recording site.

### Spike Sorting

Extracellular signals were band-pass filtered (0.3 – 3 kHz) to reveal unit activity. Potential extracellular spike waveforms were detected during recording by adaptive threshold crossing (Blackrock Microsystems). The 400 μs preceding and 1200 μs following each threshold crossing was stored and used for spike sorting. Manual spike sorting was performed with Plexon Offline Sorter. The first 3 principal components, peak-to-trough amplitude, and non-linear energy were the waveform features used for sorting. The initial waveform detection was deliberately liberal, such that it detected most unit activity as well as some noise fluctuations. The noise fluctuations provided an important baseline comparison for strict spike sorting. The degree to which a unit was different from noise was quantified with a multivariate ANOVA test (dependent variables included at least two of the waveform features). A unit was considered well isolated if the null hypothesis (unit and noise waveforms drawn from distributions with same mean) was rejected with p<0.001. If more than one unit was detected from a single electrode, each pair of units was also required to pass the same test. Moreover, each unit was required to have less than 1% of its inter-spike-intervals less than a 1 ms refractory period.

To compute the crosscorrelation in spike count between each pair of units recorded during ongoing activity we followed established methods (e.g. (Renart et al., 2010)). First, to obtain spike count vectors, the spike time stamps of each unit were 1) binned with 1 ms temporal resolution, 2) convolved with a Gaussian window with 50 ms width. The crosscorrelation coefficient was computed between all pairs of spike count vectors (2145 pairs for monkey 1, 780 pairs for monkey 2).

### Definition of LFP Avalanches

As established previously (Shew et al., 2009), we first detected negative LFP deflections (nLFPs) falling below a threshold of −3.5 SD of ongoing fluctuations *in vivo* and −6 SD of noise *in vitro*. Unlike *in vivo*, periods of quiescence between population events were clear in the *in vitro* recordings and used to define the noise baseline. nLFPs were found to occur in clusters and their sizes distributed according to a power law, the hallmark of neuronal avalanches. Two consecutive nLFPs (on any electrode) belonged to the same avalanche if the time interval between them was smaller than a threshold τ, which was determined using the probability distribution of inter-nLFP time intervals (Beggs and Plenz, 2003). We also repeated our analysis for different nLFP detection thresholds *in vivo*: 2.5, 3, 3.5, 4.25, and 5 SD, which has previously been demonstrated to not affect the power law behaviour in avalanche size distribution (Petermann et al., 2009). Our main findings were unchanged (see also next section). A complete scaling analysis of LFP avalanches for these two monkeys can be found in a recent publication (Miller et al., 2019).

### Definition of Avalanche Families

First, each spatiotemporal avalanche was represented as a binary spatial pattern with one bit per MEA electrode (Yu et al., 2011). Bits were set to 1 if the corresponding electrode recorded an nLFP during the event and otherwise set to 0. Next, patterns which included only one active site were excluded to minimize the potential inclusion of noise events. Then we sorted the events into families with similar binary patterns. K-means sorting in Matlab (Mathworks) was employed with randomly chosen seed patterns and a Euclidean distance metric. The number *k* of families to search for was decided based on the number N of population events being sorted 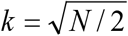 (Sánchez et al., 1979). Our objectives and the results of the k-means sorting were (1) to establish a number of families, within which events had similar spatial patterns of activation and (2) to identify a comparable number of families with relatively large spatial extent for both *in vivo* and *in vitro* data to facilitate comparison between the two approaches. This is of particular importance as typically the number of *in vitro* LFP avalanches was at the order of ~10^2^ smaller than *in vivo*. In our view, there is no single “correct” choice of *k* for experimentally recorded cortical population events, which are not likely to ever repeat exactly.

Practically, as k is reduced, families include more population events and units become less selective for families. We quantified this trend by computing the ratio of the PETH peak height *H*_*f*_ of selective unit-family pairs to the nLFP-triggered PETH peak height *H*_*nf*_, which disregards families. We found that for k = 30, 20, 10, 5, 4, 3, the ratio *H*_*f*_ /*H*_*nf*_ = 4.8, 4.4, 3.4, 3.2, 2.2, 1.0 respectively (this analysis was done for monkey 1). Note that 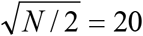 for monkey 1. Our main conclusion that some extracellular units are preferentially active for some avalanche families are not qualitatively affected by changes in k between 10 and 30. Higher values of k tended to reduce the number of events in each family, resulting in poor statistics. Moreover, using hierarchical clustering as reported previously (Beggs and Plenz, 2004;Stewart and Plenz, 2006) did not significantly change our results (data not shown). As noted in the previous section, we repeated the analysis with different nLFP thresholds. We found that for thresholds of 2.5, 3, 3.5, 4.25, and 5 SD, the ratio *H*_*f*_ /*H*_*nf*_ = 2.6, 3.5, 4.4, 6.0, 4.7 respectively (monkey 1). Thus, for all thresholds, we found a greater than 100% increase in selectivity compared to nLFP-triggered PETHs which disregard families of population events.

### Family-triggered Peri-event Time Histograms

To characterize the relationship between every unit and every avalanche family, we computed family-triggered peri-event time histograms (PETHs). If the family was comprised of *N* avalanches, then the trigger times for the PETH were the N timestamps of the first nLFPs in each avalanche. The PETHs included the 750 ms time periods preceding and following the trigger times. The bins were 50 ms in width. A PETH peak was deemed ‘selective’ if two conservative criteria were met. First, the integrated spike rate within a ±200 ms interval around the trigger time must be 3 times larger than the baseline spike rate computed in the two intervals −750 to −200 ms and 200 to 750 ms, relative to the trigger time. This criterion effectively reduces false positives but may classify units with very broad PETH peaks as non-selective. Second, the spike count in the ±200 ms interval around the trigger time must occur with probability less than 0.01 assuming Poisson spike generation of the neuron with rate λ. The rate λ was the mean spike rate calculated during the ±10 s intervals around the trigger times. The second criterion greatly reduces false positives for neurons with low firing rates, which can be common. As discussed in the main text, the number of expected false positives using these criteria was fewer than five times less than the observed number of selective unit-family pairs. To assess the delay *t* and width σ of significant PETH peaks, we fit the PETH with a four parameter Gaussian function: 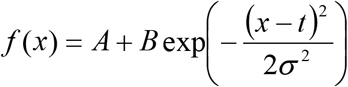.

The fit parameters *A*, *B*, *t*, and σ were determined by minimizing the summed squared differences between spike counts in each bin and the fit function. The minimization was performed with a simplex search method (Matlab function – *fminsearch*).

### Acute Slice Preparation and Recording Media

Coronal slices from medial prefrontal cortex (mPFC) or somatosensory cortex of Sprague Dawley rats were cut at 400 μm thickness (VT1000S; Leica Microsystems, GmbH) in chilled artificial cerebral spinal fluid (ACSF). We used two different types of ACSF during recordings, one for each of the two different age groups, young and old, of rats in this study. In the first protocol, referred to as the *DA/NMDA protocol* in the main text (Beggs and Plenz, 2003;Stewart and Plenz, 2006), slices were cut in ACSF saturated with 95% O_2_ and 5% CO_2_ (310 ± 5 mOsm) containing (in mM) 205 sucrose, 0.5 CaCl_2_, 7 MgSO_4_, 3.5 KCl, 26.2 NaHCO_3_, 0.3 NaH_2_PO_4_, 10 D-glucose. Prior to recording, slices were stored submerged at room temperature in ACSF containing (in mM) 124 NaCl, 1.2 CaCl_2_, 1 MgSO_4_, 3.5 KCl, 26.2 NaHCO_3_, 0.3 NaH_2_PO_4_, and 10 D-glucose. Recording was done in the same ACSF as used for storage, but with bath-application of 30 μM of Dopamine (Sigma) and 5 μM of NMDA (Sigma). In the second protocol, referred to as the *normal ACSF protocol* in the main text, we recorded under normal ACSF perfusion, but stored slices in a modified ACSF prior to recording (Shew et al., 2010). In the modified storage ACSF, Na was replaced with Choline and spontaneous population activity arises without further pharmacological manipulation when switching to the recording ACSF. Moreover, this protocol may provide a practical in vitro model for the study cortical regions that lack prominent dopaminergic projections, which may be required for the DA/NMDA protocol. For this protocol, slices were cut in modified ACSF containing (in mM) 124 choline-Cl (Sigma), 1.2 CaCl_2_, 1 MgSO_4_, 3.5 KCl, 0.3 NaH_2_PO_4_, 26.2 NaHCO_3_, 10 D-glucose, and saturated with 95% O_2_ and 5% CO_2_ (310 ± 5 mOsm). Note that choline replaces the sodium of normal ACSF. The slices were stored submerged at room temperature in the same modified ACSF as used for slicing. MEA recordings were performed in normal ACSF. All recordings were performed with ACSF saturated with 95% O_2_ and 5% CO_2,_ perfused at 3 – 4 ml/min at 35.5 ± 0.5°C. Disinhibited activity was recorded during bath-application of picrotoxin (50 μM, Sigma) in either ACSF_DA/NMDA_ or ACSF_normal_.

### Recording LFP avalanches In Vitro

Spontaneous LFP activity was recorded with integrated planar microelectrode arrays (Multichannel Systems; GmbH) that contained 59 electrodes arranged on an 8×8 grid with inter-electrode spacing of 200 μm and 30 μm electrode diameter (four corner electrodes and one ground electrode missing). Extracellular signals were recorded with 1 kHz sample rate and low-pass filtered between 1 – 200 Hz to obtain the LFP. Activity was recorded for 20 – 45 min. Experiments with fewer than 100 nLFPs were not included in our analysis. Avalanches and avalanche families were defined as described above for the *in vivo* recordings.

### Whole-cell Patch Recordings

The intracellular patch solution contained (in mM) 132 K-gluconate, 6 KCl, 8 NaCl, 10 HEPES, 0.2 EGTA, 2.2 Mg-ATP, 0.39 Na-GTP (Sigma-Aldrich). The pH was adjusted to 7.2 – 7.4 with KOH. The final osmolarity of the pipette solution was 290 ± 10 mOsm. Biocytin hydrochloride (.3%; Sigma-Aldrich) was added to the pipette for use in post fixation (4% paraformaldehyde) anatomical reconstruction. Putative pyramidal cell ~100 μm from the slice surface were visually identified by prominent apical dendritic branches and later confirmed by reconstructed morphology and/or electrophysiology. Intracellular membrane potentials were recorded in current-clamp mode (Axopatch 200B, Axon Instruments, USA), pre-amplified and low-pass filtered at 10 kHz (Cyberamp380, Axon Instruments), and digitized at 25 kHz for voltage and 5 kHz for current using the CED 1401 (Cambridge Electronic Design, UK). Data were collected with Spike2 (CED) and analysed off-line. Neurons were included in the analysis if their membrane potential was stable below −60 mV and if their action potential half width was <2.5 ms. (See Table 2 for presentation of more electrophysiological parameters.) To visualize the morphology of patched cells (e.g. Figure 6B, left), a subset of slices (n = 9) were post-processed with streptavidin-conjugated Texas Red (Molecular Probes, Inc), imaged with a Zeiss LSM 510 confocal microscope, and were stitched, projected and traced offline using Fiji ImageJ (http://pacific.mpi-cbg.de/).

## Results

### Extracellular Units Selectively Correlate with Avalanche Families in Awake Nonhuman Primates

We first studied the relationship between population avalanche patterns and single neuron activity in ongoing activity of nonhuman primates. Recordings of the local field potential (LFP; 1 – 100 Hz) were performed with high-density multielectrode arrays chronically implanted towards superficial layers of premotor cortex over the arm representation region in two macaque monkeys (Figure 1A). During the 30 min recordings, the monkeys were awake, but not engaged in any particular task. Spike sorting was used to identify 66 and 51 well distinguished extracellular units in monkeys 1 and 2, respectively (for details see Material and Methods). The firing rates of the units were 3.6 ± 9.4 Hz (mean ± SD) ranging from 0.03 to 52 Hz. The trough-to-peak time difference of unit waveforms was 345 ± 140 μs. We found average pairwise spike correlation coefficients of 0.050 ± 0.002 and 0.015 ± 0.001 for our monkeys 1 and 2, respectively consistent with previous reports (e.g. (Ecker et al., 2010;Renart et al., 2010)). In line with previous studies (Gray and Singer, 1989;Murthy and Fetz, 1996;Destexhe et al., 1999;Pesaran et al., 2002;Nauhaus et al., 2009;Petermann et al., 2009;Kelly et al., 2010;Okun et al., 2010), we observed a tendency for units to coincide with negative excursions in the LFP (Figure 1A). This was quantified by computing spike rate as a function of LFP amplitude recorded within a 50 ms windows at the same site. As shown in Figure 1B, which displays the average over all units and all times, rate increases with negative LFP amplitude as reported previously for high-density arrays based on tungsten electrodes (Shew et al., 2009).

Having demonstrated that the LFP and extracellular units are related at individual electrodes, we next explored traditional spike-triggered and LFP-triggered relationships for our recordings to identify spatial selectivity in the LFP or unit activity with respect to activity on the array. The example in Figure 1C and D, in which the location of the trigger unit is marked by the red triangle, draws attention to the spatially widespread, seemingly non-selective average nLFP activity related to local spiking. When slow LFP fluctuations were included in the analysis, i.e. band-pass filtering between 1 – 100 Hz, we found that the spike-triggered LFP waveform exhibited a broad (~0.5 sec) negative deflection with a minimum close to the trigger time (Figure 1C). When very slow fluctuations were excluded, by band-pass filtering the LFP between 3 – 100 Hz, the spike-triggered average LFP waveform displayed a sharp (~20 ms) negative peak with largest amplitude at the recording site nearest the triggering unit, in line with previous studies (Nauhaus et al., 2009;Petermann et al., 2009), yet it systematically decayed with distance from recorded unit (Figure 1D). This suggests a rather non-selective spatial relationship between low-frequency components in the LFP and single-unit activity. Next, we computed LFP-triggered averages of unit activity using the times of large negative peaks in the LFP (nLFPs) for triggers (Figure 1E). We considered all nLFPs that fell below –3.5 standard deviations (SD). For this and the remainder of the analysis in this paper, we studied the 1 – 100 Hz frequency band of LFP signals. Consistent with previous studies in awake animals (Destexhe et al., 1999;Petermann et al., 2009), we found that peri-event time histograms (PETHs) of unit counts often indicated peak firing centered on the nLFP times. Consistent with Figure 1C, units that were distant from the nLFP recording site displayed a PETH peak that was comparable with that of nearby units, on average.

Figure 1 demonstrates that, on average, the spiking activity of single neurons is related to the LFP signal. However, the spike-triggered average LFP waveform for the average unit peaks around 1 – 10 μV (Figures 1C, D), which is much smaller than the 100s of μV moment-to-moment fluctuations in the LFP (standard deviation over all electrodes was 35 ± 5 μV). Similarly, the nLFP-triggered spike histogram revealed an average increase in firing of less than 1 Hz (Figure 1E), which is a small change relative to ongoing 100s of Hz fluctuations in spike rate. The coefficient of variation for the inter-spike-interval (ISI) distributions was 2.2 ± 0.5 and the standard deviation of instantaneous spike rates (1/ISI) was 75 ± 35 Hz. These observations raise the question to what extent do average relationships faithfully represent the moment-to-moment relationships between spiking and LFP signals?

The analysis that follows was designed to answer these questions and consisted of three main steps. First, we identified neuronal population events based on the spatial patterns of LFP signals afforded by multi-site recordings. Second, we sorted the population events into ‘families’ of like events, based on which sites exhibited negative LFP deflections during each event. Third, we tested each unit individually for family-specific changes in firing. If our hypothesis is correct, we should find that certain units fire selectively during certain families, while other units prefer other families.

Our definition of a population event is motivated by two observations: 1) nLFPs are associated with increased spiking activity and 2) LFP signals recorded simultaneously from different sites are often highly correlated (e.g. Figures 1A and 2A, (Destexhe et al., 1999;Leopold et al., 2003;Nauhaus et al., 2009). Therefore, we define a population event to be a set of nLFPs (typically from many recording sites), which occur together sufficiently close in time. Specifically, if the time interval between two consecutive nLFPs is less than a threshold τ, we assign them to the same population event. The threshold τ is chosen based on the inter-nLFP-interval distribution, which was bimodal; τ is between the peaks in the distribution, thus distinguishing the long time-scale which separates events and the short time-scale activity in nonhuman primates (Leopold et al., 2003;Miller et al., 2019), slow timescale dynamics were dominant, although monkey 1 did show a slight increase in gamma band power near 30 Hz compared to 10 – 20 Hz range (Figure 2B). We recorded 1,308 and 2,016 population events for monkeys 1 and 2, respectively. Population events were diverse in spatial extent, spanning 9.6 ± 16.5 and 9.4 ± 16.2 electrodes (mean ± SD) for monkeys 1 and 2. We defined the size of a population event as the summed amplitudes of all the nLFPs comprising the event and demonstrate that the distribution of the population event sizes was close to a power-law with exponent −1.5 (Figure 2C). This power-law event size distribution indicates that the dynamics we study here are ‘neuronal avalanches’ (Beggs and Plenz, 2003), in line with previous studies of ongoing activity in awake monkey cortex (Petermann et al., 2009;Klaus et al., 2011;Miller et al., 2019). Next, for each avalanche, we generated a representative binary 10×10 pixel pattern (corner electrodes missing), which indicates which sites were active during the event (1=active, 0=inactive; (Yu et al., 2011)). Figure 2A exemplifies 3 s of simultaneous LFP recordings with an avalanche occurring about 1.5 s into the example (red dots mark nLFPs). The upper left in Figure 2D shows the corresponding binary pattern for this occurrence. Next, we used a k-means algorithm to find families of avalanches with similar activation patterns (Beggs and Plenz, 2004;Stewart and Plenz, 2006). Four example patterns from one family are shown in Figure 2D. The nLFP raster in Figure 2E shows all nLFP times and sites during a 30 min recording from monkey 1. Figure 2F displays the corresponding binary patterns derived from the nLFP raster sorted into families of similar patterns. The occurrence-times of events in one family were typically scattered throughout the 30 min recording. We note that k-means sorting also results in one ‘misfit’ family comprised of many small and local events that repeated rarely during the recording and will not be considered further for this analysis (e.g. family *a* in Figure 2F). The sorted raster of binary patterns for monkey 2 is shown in Figure 2G.

**Figure 2.**
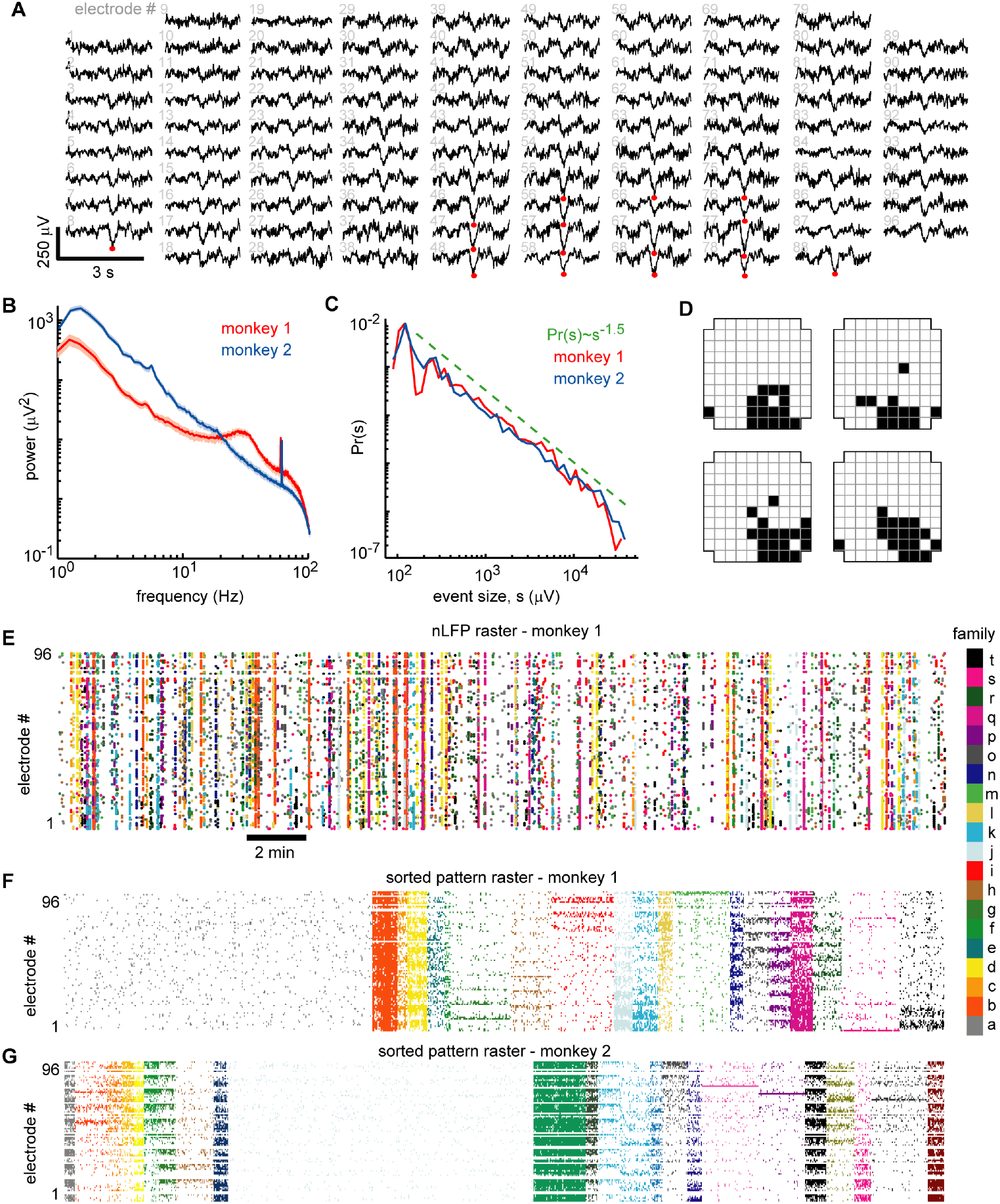
Ongoing neuronal avalanches are composed of repeating spatial nLFP patterns. **A.** Displayed are 3 s LFP recordings arranged to match the spatial layout of the 96 recordings sites. Multiple nLFPs (red dots) often occurred together within ~100 ms across multiple sites – we defined such occurrences as ‘population events’ (see ‘Experimental Procedures’). **B.** Power spectra of LFP were broadband showing that low frequency fluctuations dominate the signal. A prominent ~30 Hz oscillation was present in monkey 1 (see also (Miller et al., 2019)). All recording sites were analyzed – median (line) and lower to upper quartile (shaded region) are shown. **C.** Distributions of population event sizes *s* demonstrate that the activity is neuronal avalanches, defined by *Pr(s)*^~^*s*^−*1.5*^ (Beggs and Plenz, 2003;Petermann et al., 2009). **D.** The spatial locations of nLFP avalanches were represented with a binary pattern (Yu et al., 2011). The upper left pattern corresponds to population event in A. The other three patterns were similar but occurred at different times and are shown as typical like-examples extracted by our algorithm use. **E.** Raster of all nLFP times and locations. Vertical clusters of nLFPs with matching color belong to one avalanche. **F.** Avalanches were sorted into families with like patterns. The sorted avalanche raster of monkey 1 is shown. Color code identifies family and is same as in E. Note that family *a* (gray, left) is comprised of typically small avalanches that were not similar to many others. **G.** Sorted avalanche raster of monkey 2.

With the population events sorted into families, the next step was to determine whether units fired selectively for families. To accomplish this, we computed family-triggered spike rate PETHs. One PETH was computed for each unit-family pair, triggered on the times of the first nLFP in each population event of the family. Figure 3 shows examples from five units and a subset of families from monkey 1. Figure 3A shows the average pattern for each example family. Figures 3B – F (top) show family-triggered PETHs from the five example units, whose locations are color-coded on the grid in Figure 3A (right). Below each PETH, we display the individual family-triggered spike trains from which the PETH is constructed (Figures 3B – F, bottom). Our main finding was that extracellular units were reliably and selectively active for avalanche families identified in the LFP. Some units were reliably active during multiple families (n=32 and 4 for monkeys 1 and 2 respectively; e.g. families *o* and *d* in Figure 3B – D), while other units (Figure 3E, F) fired reliably for only one family (n=13 and 19), or none at all (n=21 and 28). A summary of average patterns for all families of monkey 1, the grand average over all population events, disregarding families, and spatial location of all units recorded are shown in Figure 4. All units and their respective family selectivity are summarized in Figure 5A for both monkeys. In monkey 1, we found 124 selective unit-family pairs with a strong change in firing revealed by the family-triggered PETH. In monkey 2, we found 29 selective unit-family pairs. Here, we adopt a conservative definition of ‘selective’, requiring a strong increase or decrease in firing compared to baseline (see Experimental Procedures for detailed definition). In both monkeys, the number of strong relationships was more than five times greater than the number expected by chance (9 and 5 for monkeys 1 and 2). To demonstrate this, we repeated our analysis with randomized spike times – each time was shifted by a random amount between 1 and 10 s.

**Figure 3.**
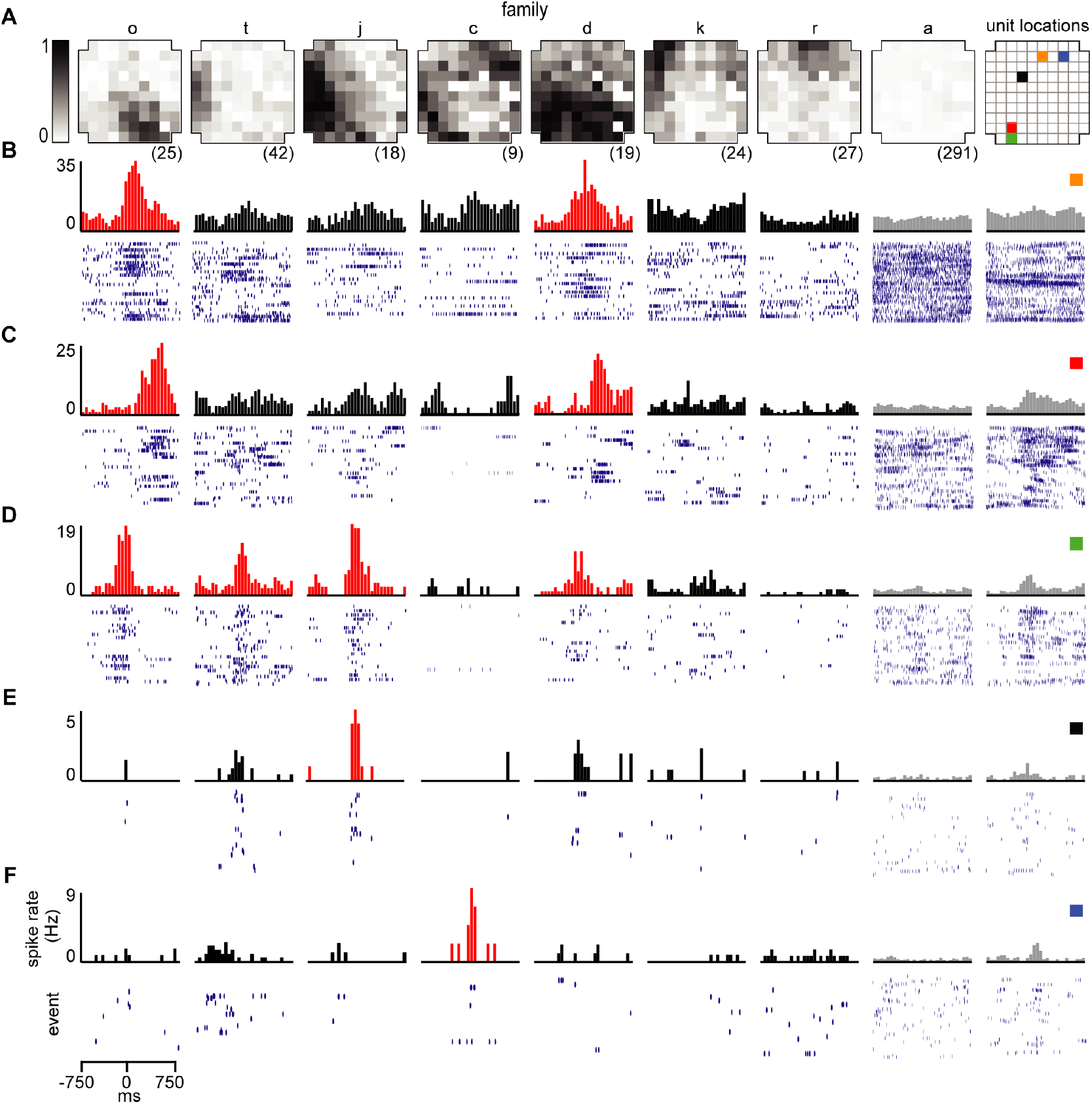
Reliable ensembles of spiking units underlie avalanche families. **A**. The average pattern for 8 example avalanche families. Letter family labels correspond to Figure 2E, F of monkey 1. Grayscale indicates fraction of events in which the site participated. *Right*: color code for locations of units in *B* – *F*. **B – F**. Each column displays family-triggered spike rate PETHs (top) and spike rasters (bottom) for five example units. For example, the left-most PETH and spike raster in *B* indicates that this unit increases its firing rate selectively during the occurrence of family o. The temporal bin width for the PETHs was 50 ms. The right-most column includes PETHs and rasters triggered on all nLFPs recorded at the same site as the unit, disregarding families.

**Figure 4.**
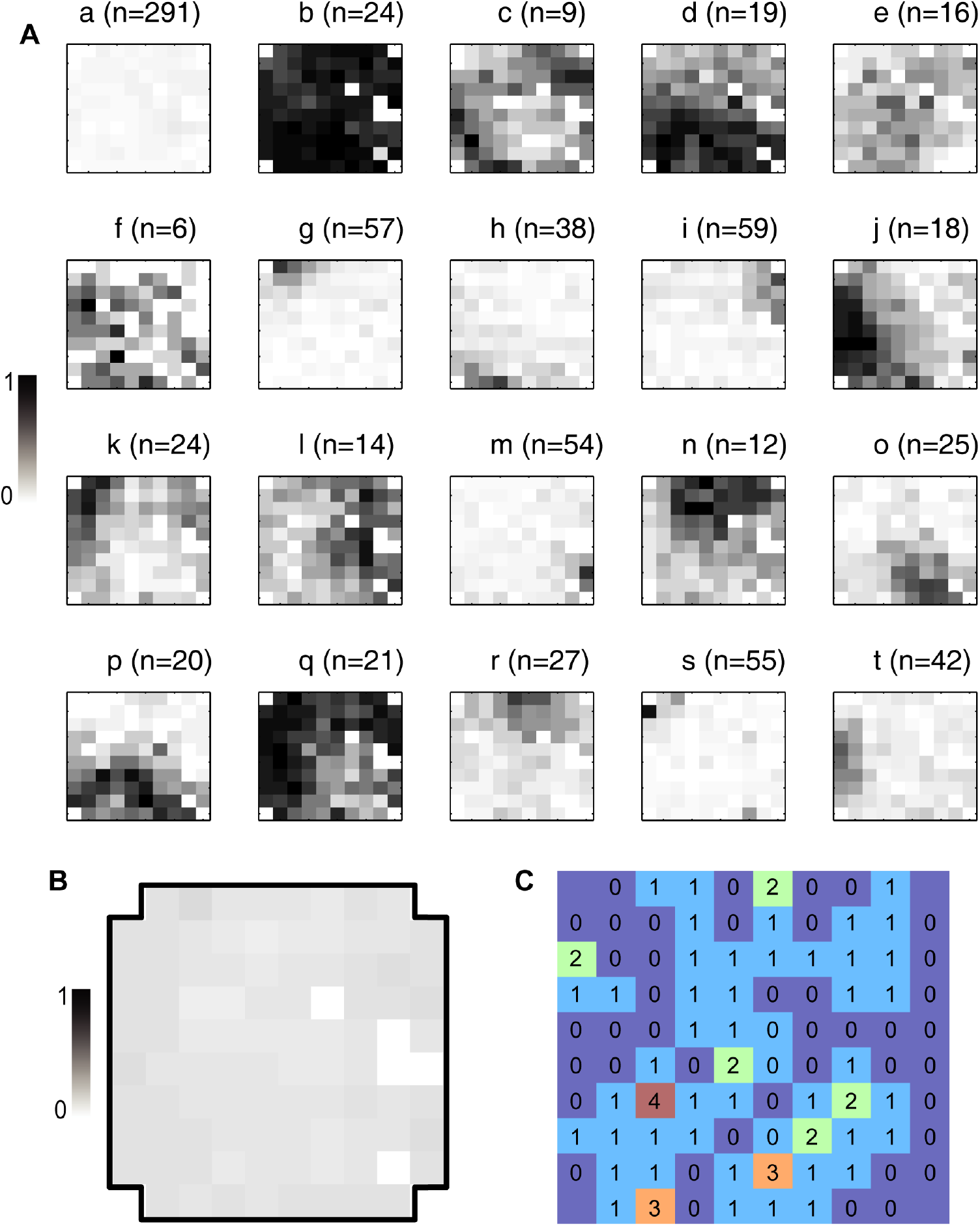
Pattern and unit overview for monkey 1. **A.** Average activation patterns from the full set of avalanche families for monkey 1 of which a subset of families is shown in figure 3A. Grayscale indicates the fraction of population events within the family during which the site was active. The letter indicates the family as labeled in figure 3 and n indicates the number of avalanche occurrences comprising the family. **B.** Grand average activation pattern over all avalanches. Grayscale indicates fraction of population events during which an nLFP was recorded. Note that the average family patterns in figure 3A are not well represented by the full average. **C.** The number of units recorded at each site. This can be compared with figure 3E, F to see what fraction of units were selective for the example patterns.

**Figure 5.**
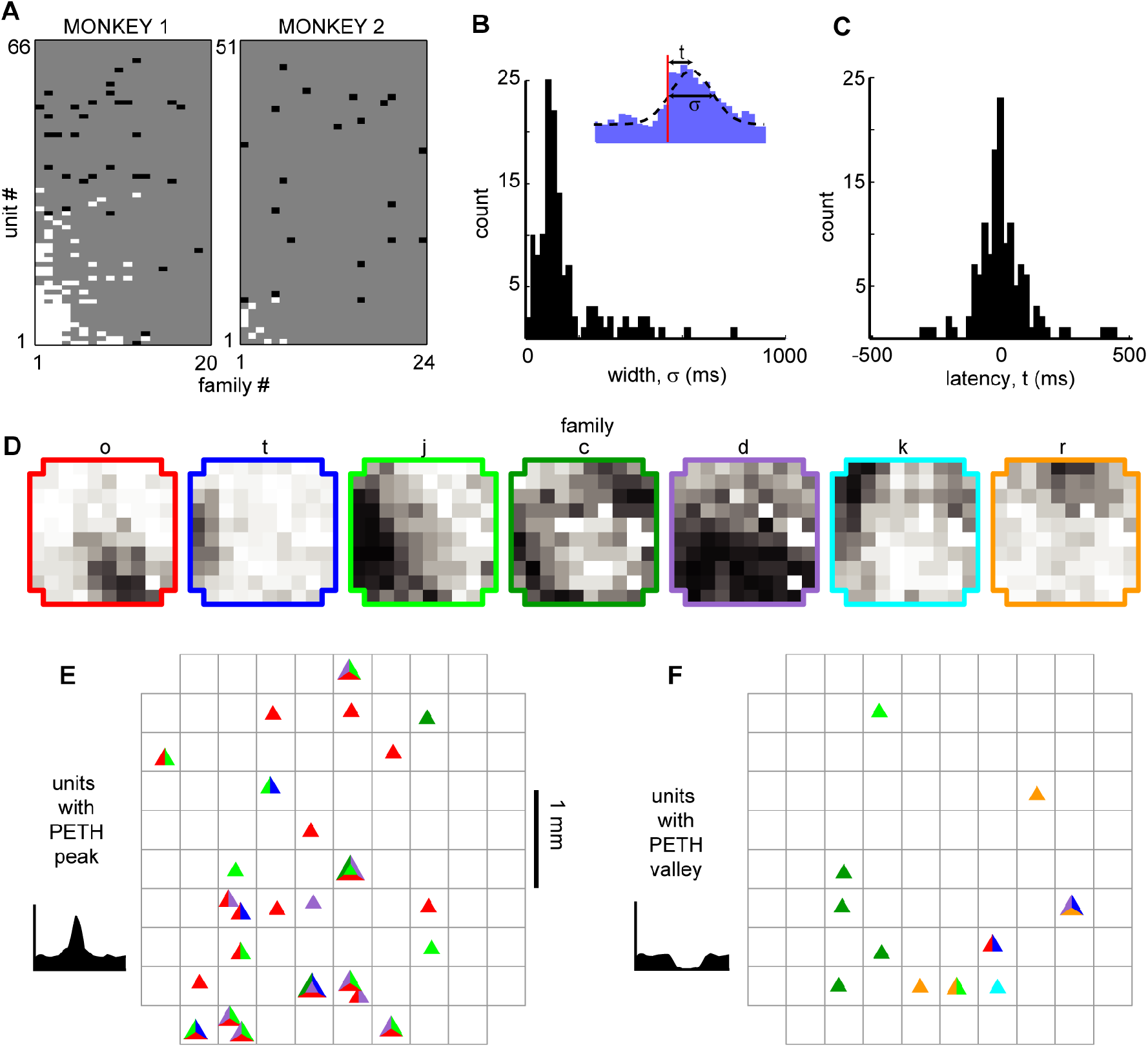
Avalanche families recruit temporally and spatially diverse unit ensembles. **A.** Summary of all unit-family relationships for both monkeys. White pixels indicate firing increase, black pixels indicate decrease in firing. **B.** We estimated the latency *t* to peak firing and the width σ of the PETH peak for each strong unit-family pair, by fitting a Gaussian function with an offset (*inset*). The distribution in PETH widths peaked around 100 ms but was also widespread indicating that the temporal precision of unit-family relationships was variable. **C.** Similarly, peak unit firing centered near t = 0 but could precede or follow the occurrence of a family by up to 100 ms. **D.** Color-code for same average family patterns as shown in Figure 3. **E.** Displayed are the locations of units which selectively increased their firing for the families in D. Each unit is represented with a triangle. The colors within each triangle indicate which families the unit was selective for. **F.** Same as in E, but for units that selectively decreased their firing.

Among the units and families that were strongly related, we found that the temporal precision of unit participation in families was varied. For example, Figures 3B, C have PETH peaks that are much broader than those in Figures 3E, F. Moreover, the latency from trigger time to PETH peak also varied.

To quantify the width and latency of the PETH peaks, we fit a Gaussian function to the PETH (Figure 5B, inset, Experimental Procedures). We found that the latencies were broadly distributed 5 ± 107 ms (mean ± SD) and 0.4 ± 106 ms for monkeys 1 and 2 respectively (Figure 5B). The PETH peak widths, i.e. standard deviation parameter of the Gaussian fit, were 140 ± 136 ms and 170 ± 110 ms for monkeys 1 and 2 respectively (Figure 5C).

The locations of the units which fired reliably during a given family were distributed diffusely over the majority of the 4 mm × 4 mm recording region. The unit locations often did not overlap with the location of nLFPs that comprised the pattern (e.g. Figure 3B, family *o*). Figures 5D – F summarizes the locations of all units that reliably changed their firing rate for the set of example families shown in Figure 3 for monkey 1. Figures 5E and F display locations of units which showed a reliable increase and decrease in firing rate, respectively.

The grand average nLFP-triggered spike histograms and spike-triggered average LFP shown in Figure 1 conceal the richness of the relationship between different units and different families of LFP population events. For example, comparing the nLFP-triggered spike histograms in Figure 1E to the family-triggered PETHs in Figure 3, we see that family-triggered PETHS often had much larger or sharper peaks. This can also be seen by comparing the family-triggered PETHs to the rightmost PETHs in Figure 3, which were triggered on the times of all nLFPs that occurred on the electrode which recorded the unit. Quantitatively, we found that the selective unit-family pairs (as defined above) exhibited a PETH peak that was 4.4 ± 8.8 times larger than the nLFP-triggered PETH peak for monkey 1 and 4.3 ± 6.9 times larger for monkey 2. These results demonstrate that if all units and population events are averaged together as in Figures 1B – E, one underestimates the strength and spatiotemporal complexity of the relationship between unit activity and the LFP.

### Synaptic Inputs to Layer 2/3 Pyramidal Neurons Selectively Occur During Avalanche Families in Rat Acute Slices

We have shown above that local ensembles of spiking neurons are closely related to LPF based avalanche patterns in the cortex of awake monkeys. We next carried out combined whole-cell patch clamp and multi-site LFP recordings in acute slices of rat somatosensory and medial prefrontal cortex (Figure 6). Since afferent fibers from distant regions are severed in the acute slice, this preparation allows us to investigate the selectivity of intrinsic dynamics in local cortical circuits. We focused on the role of layer 2/3 pyramidal neurons and carried out control experiments with pharmacologically blocked fast GABA_A_-receptor mediated synaptic inhibition.

**Figure 6.**
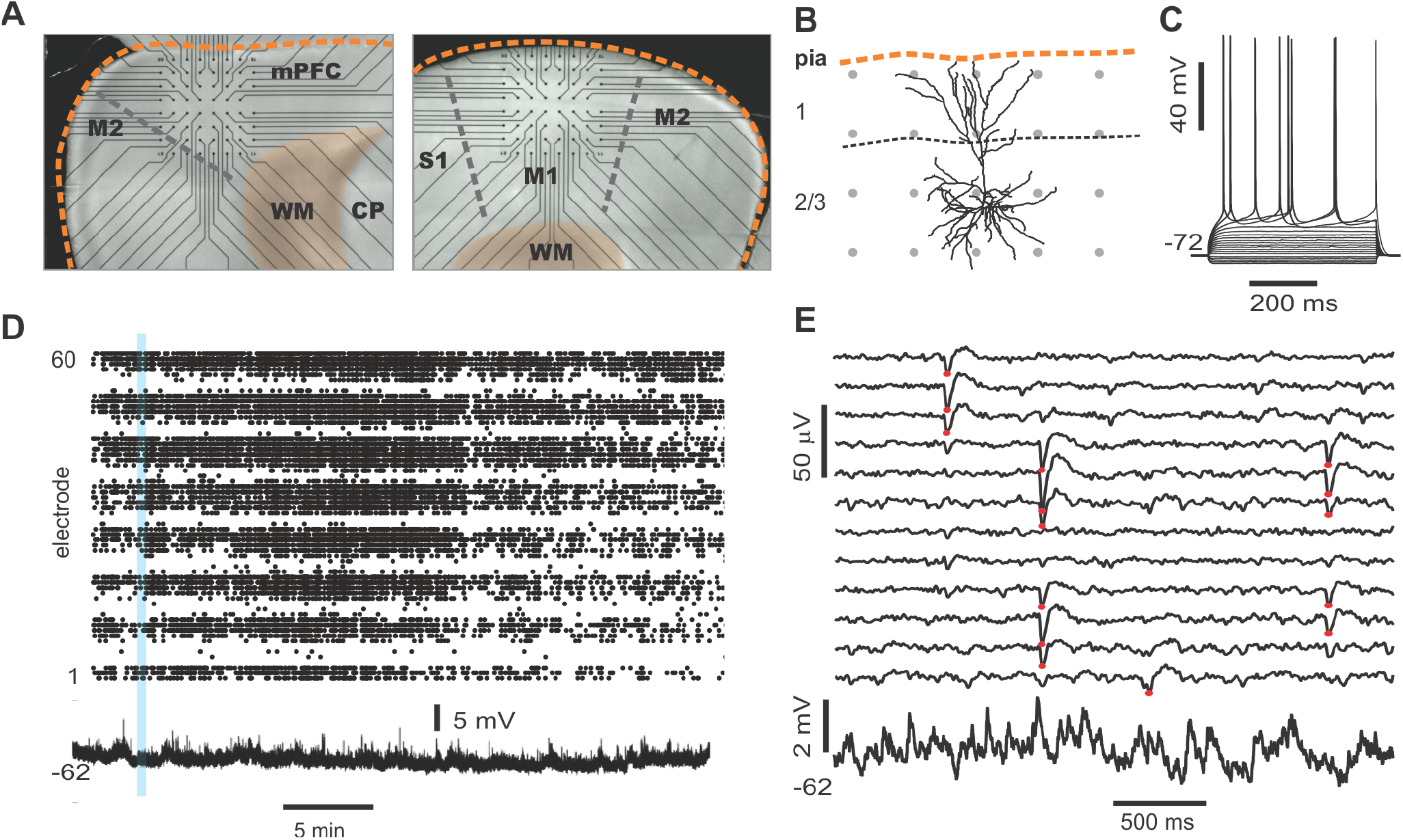
Simultaneous multi-site LFP and whole-cell patch recordings *in vitro*. **A.** Trans-illuminated pictures that display placement of acute coronal slices from rat cortex on a planar multi-electrode array MEA), visible as straight connection leads ending in recording electrodes (*black dots*). *Left*: Example for medial prefrontal cortex (mPFC) recordings. Medial cortex border is oriented upwards. *Right*: Example of somatosensory cortex (M1) recording. Dorsolateral axis is oriented upwards. *Scale*: inter-electrode distance is 200 μm. *Orange broken line*: cortical border. *WM*: white matter. *CP*: caudate putamen. **B.** Example of a reconstructed layer 2/3 pyramidal neuro (confocal image) to illuminate size relationships between single neurons and spacing of MEA electrodes (*grey dots*). **C.** Sub- and suprathreshold voltage esponses to step current injections of whole-cell patched pyramidal cell. **D.** Example of spontaneous nLFP activity on the full array (electrodes 1 – 60 ordered in groups of 8 per row) in the presence of NMDA/DA, which typically lasts for >30 min (Stewart and Plenz, 2006). The time course of the imultaneously recorded intracellular membrane potential from a whole-cell patched neuron is displayed at the bottom. **E.** Five seconds of ongoing LFP population activity recorded from a subset of MEA electrodes spanning layer 2/3 and intracellular membrane potential taken from period in *D* (*blue bar*) at higher spatiotemporal resolution. *Red dots* indicate suprathreshold nLFPs. Note absence of any apparent, straightforward relationship between single neuron activity and nLFPs.

In a first set of *in vitro* experiments, population activity was elicited in acute coronal slices from medial prefrontal cortex (mPFC) and motor cortex (M1) of adult rats (age 7 – 9 weeks) induced by continuous bath application of 30 μM Dopamine (DA) and 3 μM NMDA as reported previously (Beggs and Plenz, 2003;Stewart and Plenz, 2006). In a second set of experiments, slices were taken from mPFC and M1 of young rats (age 2 – 3 weeks) using a choline-based, protective slicing solution followed by recording spontaneous activity in normal ACSF (for details see Experimental Procedures). Multi-site LFP was recorded using planar 60-electrode MEAs covering a 1.6 × 1.6 mm^2^ region with an interelectrode distance of 200 μm (Figure 6A). There were several notable differences in basic parameters between these two protocols. Spontaneous LFP activity in ‘normal ACSF’ of young slices was about 10 times higher in rate and aggregate LFP amplitude compared to the DA/NMDA induction protocol for slices from adult rats (Table 1). We identified all cells as putative pyramidal neurons based on a combination of morphology, I/V-responses, and action potential properties (Figure 6B; Table 2). For most cells, we in addition obtained extensive measures of action potential firing, which demonstrated that the increase in LFP activity for the younger slices correlated with a significantly longer action potential width for pyramidal neurons typical for immature neurons (Table 2). Thus, the two protocols allowed for examining avalanche and single neuron activity under two largely different rates of activity. Except where noted, all of the following observations were found for both protocols. We defined neuronal avalanches as described previously (Beggs and Plenz, 2003;Stewart and Plenz, 2006) and sorted them into families exactly as in the *in vivo* data analysis. We note that, unlike our *in vivo* recordings, in which the MEA matrix was placed horizontally within layer 2/3, the *in vitro* MEA spanned multiple cortical layers across the coronal slice with the upper most row placed along the medial (mPFC) or dorsal/dorsolateral border of the cortex (M1). In line with our previous reports (Stewart and Plenz, 2006;2007;Petermann et al., 2009), we found that LFP activity occurred predominantly in layer 2/3 (Figure 7A,B) for both protocols. In line with our *in vivo* observations, these predominantly layer 2/3 nLFP patterns distributed in sizes according to a power law that was sensitive to temporal shuffling, again as show in our original paper on neuronal avalanches in the acute cortex slice (Beggs and Plenz, 2003) (Figure 7A,B).

**Table 1.** Comparison of *in vitro* population activity for the different conditions of recording (mean ± S.E.M). * P<0.05.

|  | DA/NMDA | Normal ACSF | PTX |
| --- | --- | --- | --- |
| Number of slices (n) | 85 | 42 | 9 |
| Spontaneous activity duration (s) | 1,486 ± 538 | 1,586 ± 667 | 1,306 ± 629 |
| Total nLFP count from all sites (n) | 596 ± 810 | 5,700 ± 9130 | 4,733 ± 6677 |
| Rate of nLFPs at single site (Hz) | 0.38 ± 0.46 | 3.7 ± 4.8 | 4.3 ± 5.1 |
| Integrated nLFP amplitudes from all sites (mV) | -5.8 ± 7.1 | -80 ± 113* | -158 ± 186 |

**Table 2.** Action potential Electrophysiological parameters for whole-cell patch recordings of pyramidal neurons, (mean ± S.E.M). * P<0.05. Only neurons for which reliable action potential measures were obtained are listed.

|  | DA/NMDA | ACSF | PTX |
| --- | --- | --- | --- |
| Number of cells (n) | 71 | 36 | 12 |
| Resting potential (mV) | $-71 \pm 4$ | $-68 \pm 5$ | $-71 \pm 5$ |
| Action potential threshold (mV) | $-35 \pm 5$ | $-38 \pm 4$ | $-34 \pm 6$ |
| Action potential amplitude (mV) | $88 \pm 9$ | $87 \pm 9$ | $87 \pm 7$ |
| Action potential width (ms) | $1.0 \pm 0.4$ | $1.68 \pm 0.6^*$ | $1.0 \pm 0.5$ |
| After hyper-polarization amplitude (mV) | $8.8 \pm 2.7$ | $8.2 \pm 2.5$ | $10 \pm 4$ |

**Figure 7.**
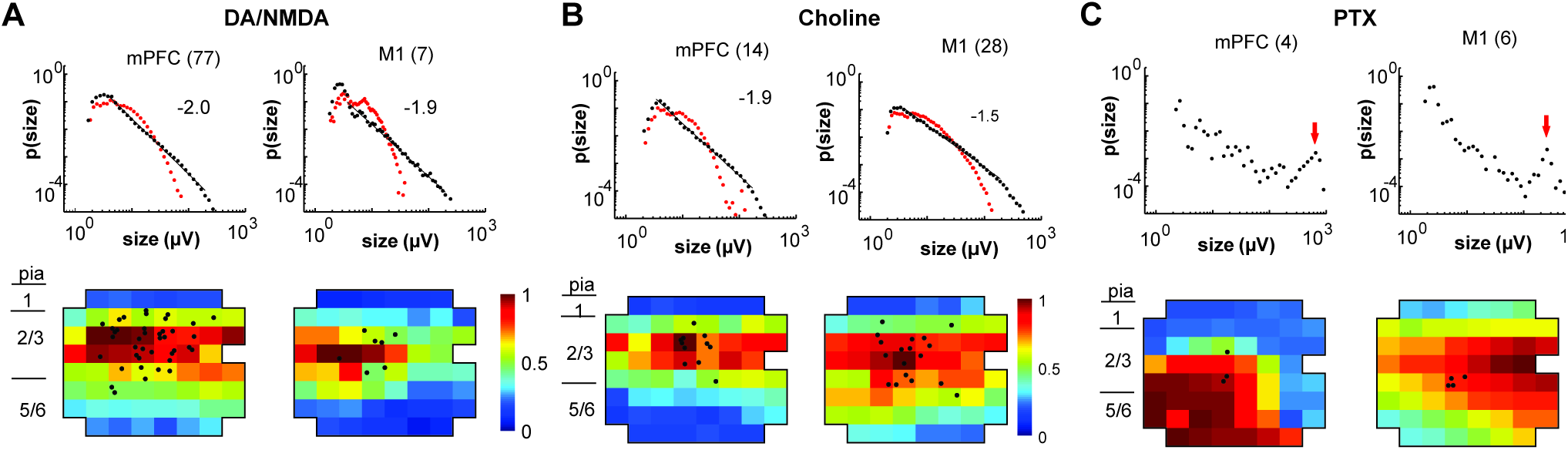
Overview of total avalanche activity and location of whole-cell patched pyramidal neurons in layer 2/3 for all recording conditions. **A.** Avalanche size distribution obtained under DA/NMDA conditions for mPFC and M1 experiments. Size distributions reveal power laws in line with neuronal avalanches, which are destroyed by shuffling (*red*). Numbers indicate total number of slice experiments for each region. Bottom density plots indicate population activity measured on the MEA aligned to medial (mPFC) and dorsal (M1) border averaged over all slice experiments. Approximate layers as a visual guide indicated on the left (MEA, 8 × 8 electrodes, 200 μm interelectrode distance, corner electrodes missing, additional ground electrode in 4^th^ row on the right;). Color indicates activation rate (averaged over experiments). Black markers indicate soma locations of patched neurons on the array. **B**. Same as in *A* for choline condition. **C**. Same as in *A* under disinhibited condition in the presence of picrotoxin (PTX). Arrow for PTX points to the predominance of system size spontaneous activations under disinhibition. Event size distributions were consistent with neuronal avalanches.

An example of a simultaneously recorded intracellular membrane potential from a layer 2/3 pyramidal neurons and the spontaneous LFP on the MEA is shown in (Figures 6B, C and 7A, B). Upon wash-in of DA/NMDA, ongoing LFP activity emerged and the intracellular membrane potential depolarized by ~4.0 ± 3.5 mV. As observed *in vivo*, LFP avalanche patterns were very diverse, but certain patterns tended to repeat during the course of a recording. Figure 8A displays an unsorted nLFP raster of avalanches. Figure 8B shows corresponding sorted raster into color-coded avalanche families. Since action potential firing was very low in the patched neurons, our goal here was to test whether neurons displayed significant subthreshold membrane potential changes with respect to particular avalanche families. To this end, we performed family-triggered averages of the membrane potential recordings.

**Figure 8.**
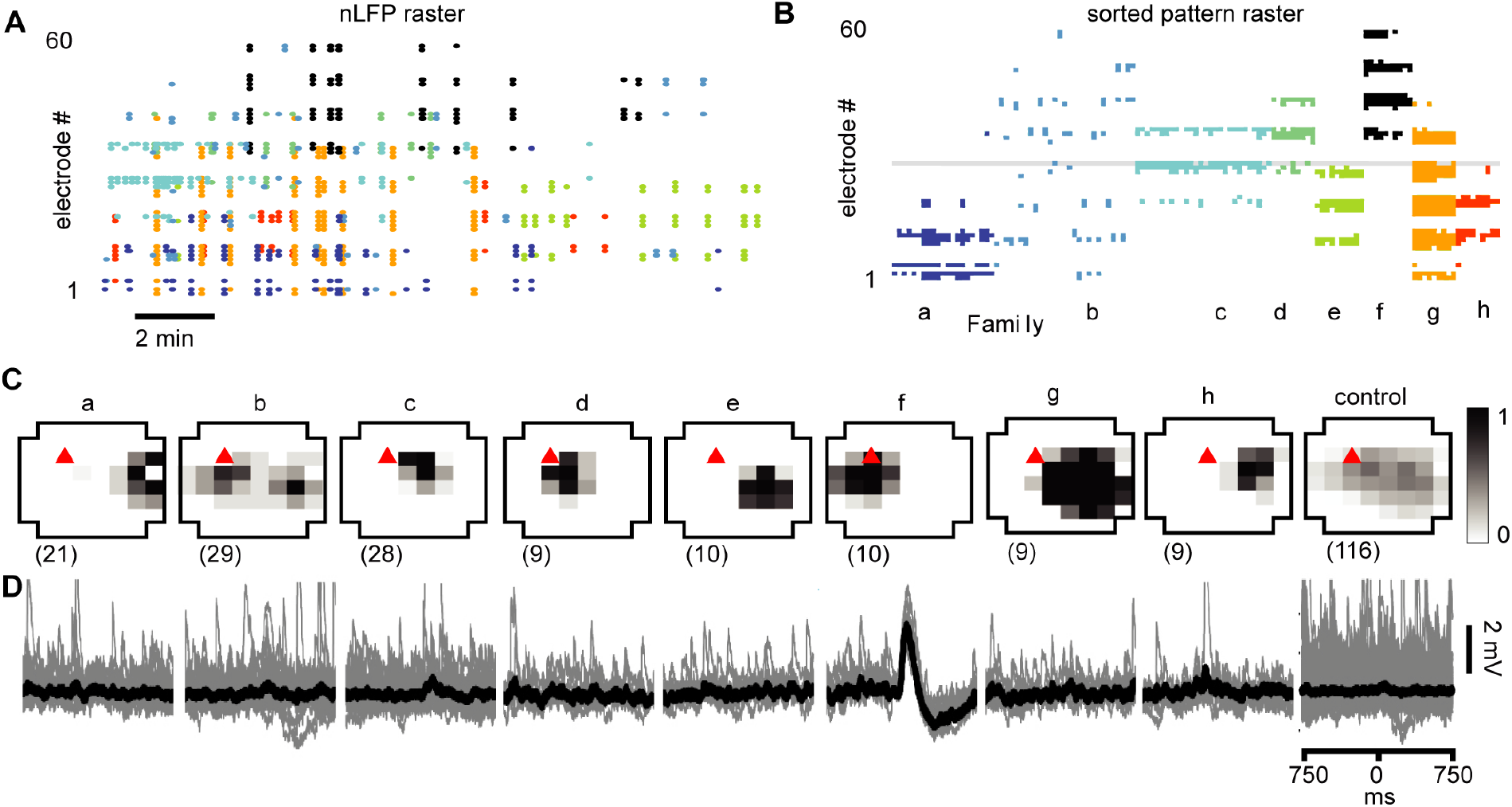
Select responses in layer 2/3 pyramidal neurons to families of NMDA/DA induced LFP avalanches. **A.** nLFP raster on the MEA (15 min recording; mPFC). **B.** Sorted nLFP avalanche raster. Color in *A* and *B* indicates avalanche family. *Gray line* indicates electrode nearest to neuron soma. **C.***Top:* Average spatial LFP pattern for all avalanche families. Grayscale indicates fraction of site participation for each family. *Red triangle* marks soma location of patched pyramidal neuron. *Control* includes all avalanches except those in family *f*. *Bottom:* Family-triggered average intracellular membrane potential time course (*black*). *Number in brackets* indicate family members. Pre-averaged individual traces shown in *gray*. Family *f* generated reliable input to the patched cell. Corresponding *control* demonstrates no reliable trigger with other family patterns outside *f*.

Our main finding from the *in vitro* recordings was that pyramidal neurons displayed reliable subthreshold membrane potential responses only from select avalanche families, in line with our *in vivo* results. Examples of family-triggered membrane potentials changes for one neuron using the DA/NMDA protocol are shown in Figure 8D below the average activity pattern for the corresponding families in Figure 8C. A corresponding example from the normal ACSF protocol for young slices is shown in Figure 9.

**Figure 9.**
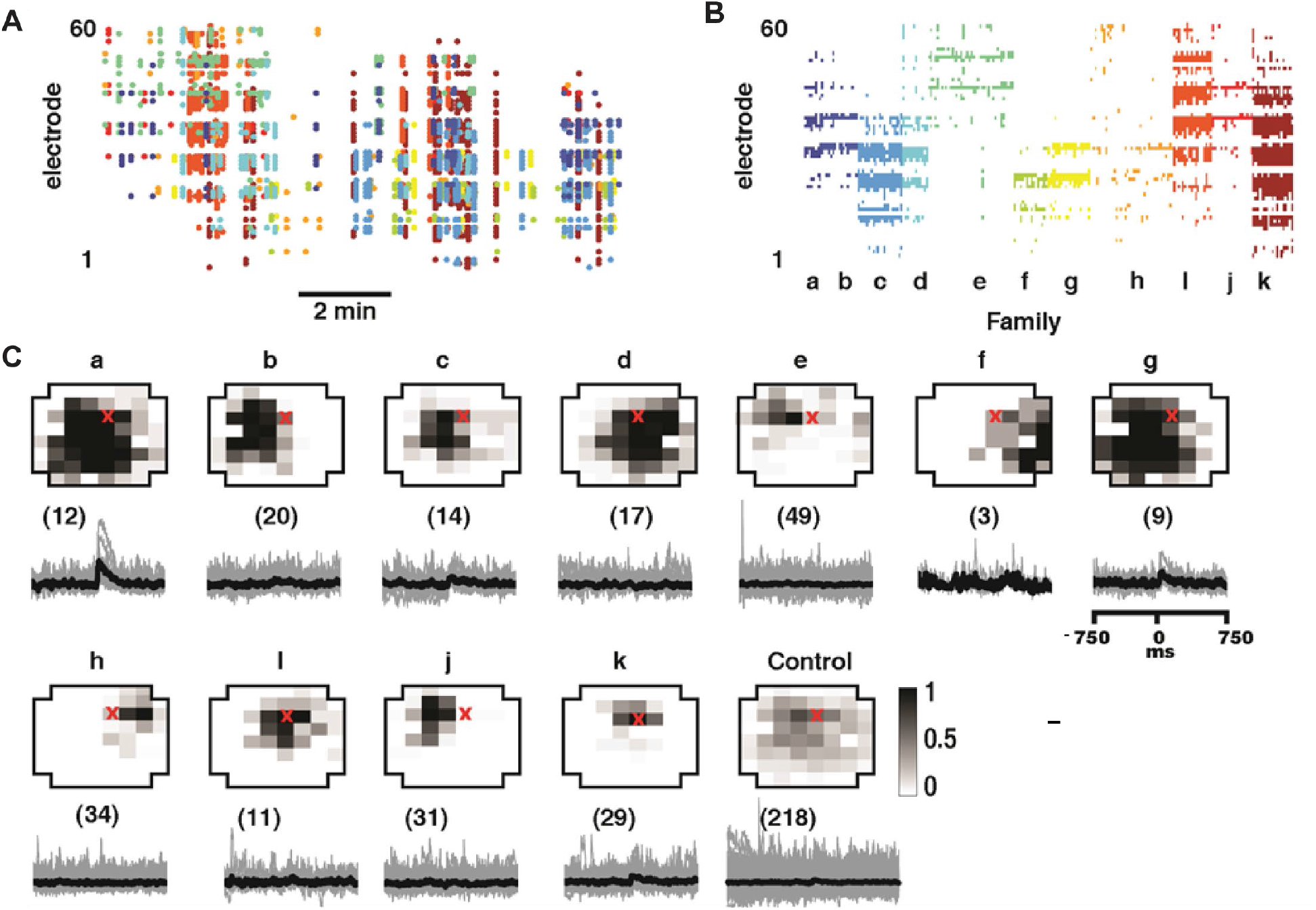
Select responses in layer 2/3 pyramidal neurons to families of LFP avalanches under normal ACSF from choline-protected young slices. **A.** nLFP raster on the MEA (10 min recording, M1). **B.** Sorted nLFP avalanche raster. Color in *A* and *B* indicates avalanche family. **C.***Top*: Average spatial LFP pattern for all avalanche families. Grayscale indicates fraction of site participation for each family. *Red triangle* marks soma location of patched pyramidal neuron. *Control* includes all avalanches except those in family *f*. *Bottom:* Family-triggered average intracellular membrane potential time course (*black*). *Number in brackets* indicate family members. Pre-averaged individual traces shown in *gray*. Note that families *i* and *k* do not reveal significant intracellular responses despite substantial spatial overlap with soma location Families *a* and *g* correlated with reliable intracellular responses in the patched pyramidal neuron. Corresponding *control* demonstrates no reliable trigger with non-*a*,*g* families.

We quantitatively identified neurons that receive significant input from avalanche families as those with family-triggered average membrane potential that exhibited a deflection during the 50 – 200 ms window following the trigger with an SD that was 3 times bigger than the SD of the pre-trigger baseline (−500 to −50 ms). Out of 84 recorded cells, only about 50% received input from just a few avalanche families (for a summary see Figure 10A, B). Thus, we conclude that participation of single neurons in avalanche dynamics was typically selective.

**Figure 10.**
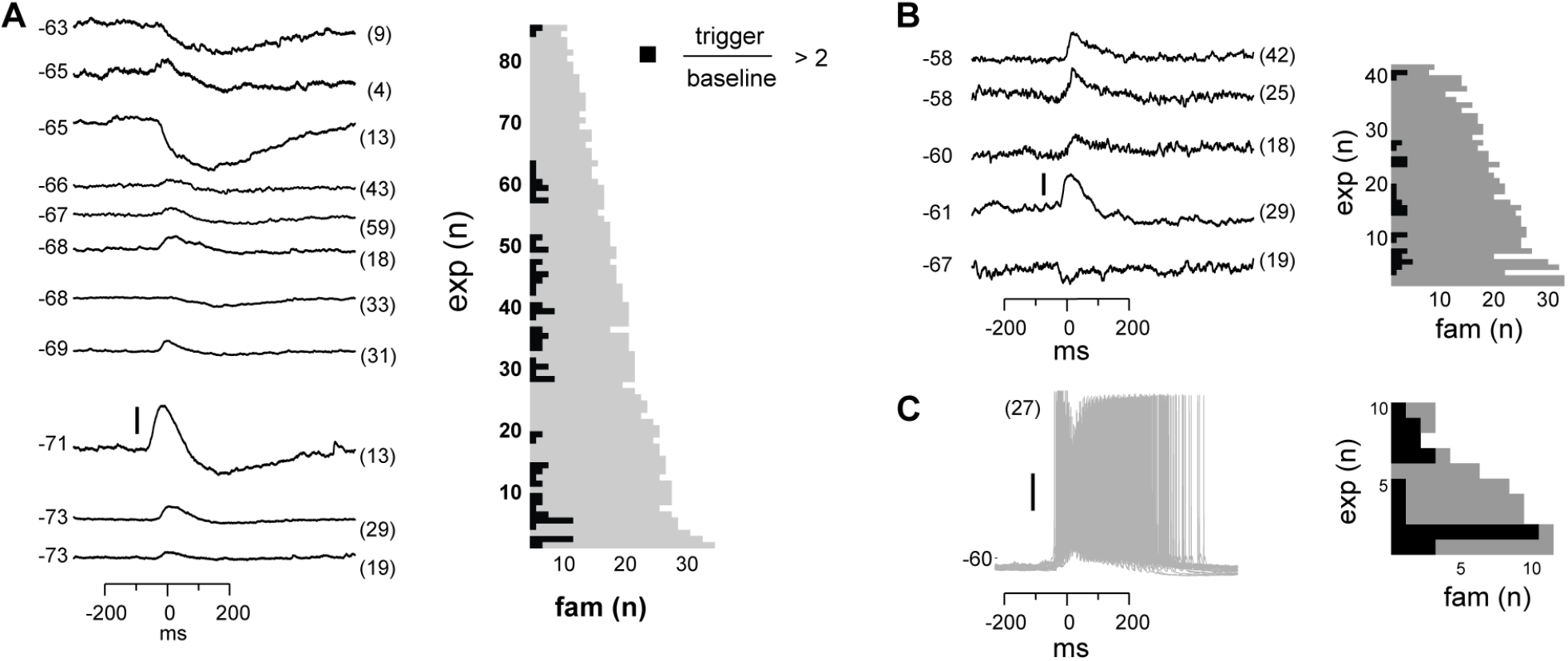
Summary of significant membrane potential responses within each experimental group. **A.** NMDA/DA condition: 42 neurons were found to receive reliable and selective input from particular avalanche families. *Left*: subset of 11 family-triggered average membrane potential time courses with large amplitude changes ranked according to trigger-preceding average membrane potential, shown at the lower right of each trace. Number of family triggers indicated by number in brackets to the right. Note that responses could be depolarizing, hyperpolarizing or a depolarization followed by a hyperpolarization. *Right*: Summary plot of total number of avalanche families and number of families with select intracellular responses. Note that most families will not trigger a response in a randomly recorded pyramidal neuron. **B.** Summary for normal ACSF condition. **C.** Summary for PTX condition.

As in our *in vivo* results, the relationship between the input to the neuron and the population activity in Figures 8 and 9 would be missed or, at best, underestimated if one averaged over all avalanches and all neurons. Here we emphasize this point by computing the average membrane potential triggered on all avalanches except the selective family (Figure 8C and 9C; *right*, ‘control’), which results in no significant average membrane deflection. These results indicate that accounting for the spatial pattern of avalanches is crucial to identify the relationships we present. In fact, LFP activity recorded with a randomly chosen single electrode from our multi-site recordings is likely to be uncorrelated to the input to any particularly patched neuron. For example, the recording site marked by the gray line in Figure 8B is within 500 μm of the soma of the patched neuron, but the events that occur at that site are not in the family the neuron is selective for. Therefore, input to the neuron would appear unrelated to the LFP signal recorded at this site.

Finally, we investigated whether the selectivity encountered in our analysis might be due to the specific algorithmic approach to avalanche identification or might be maintained dynamically by the cortical network. To this end, we examined the role of fast GABA_A_-receptor mediated synaptic inhibition. It is well established that suppression of inhibition reduces selectivity of neural response to sensory stimuli (e.g. (Kyriazi et al., 1996)). We tested whether the selectivity for particular families of ongoing LFP patterns also depends on inhibitory signaling. We bath applied GABA_A_ receptor antagonist picrotoxin (50 μM). Under such disinhibited conditions, ongoing activity was comprised of stereotyped population events with much larger LFP amplitude and spatial extent than observed with intact inhibition (Figure 11; see also Table 1). This loss in diversity of population events changes the size distribution away from avalanches displaying a prominent system-wide peak (Figure 7C, arrow). Similarly, the loss in diversity of patterns precludes selective participation of single cells; a cell cannot be selective with only one family to choose from. Intracellular recordings (see Table 2) confirmed the loss of selectivity; they revealed membrane depolarization or action potentials during nearly every population event (Figure 10, 11).

**Figure 11.**
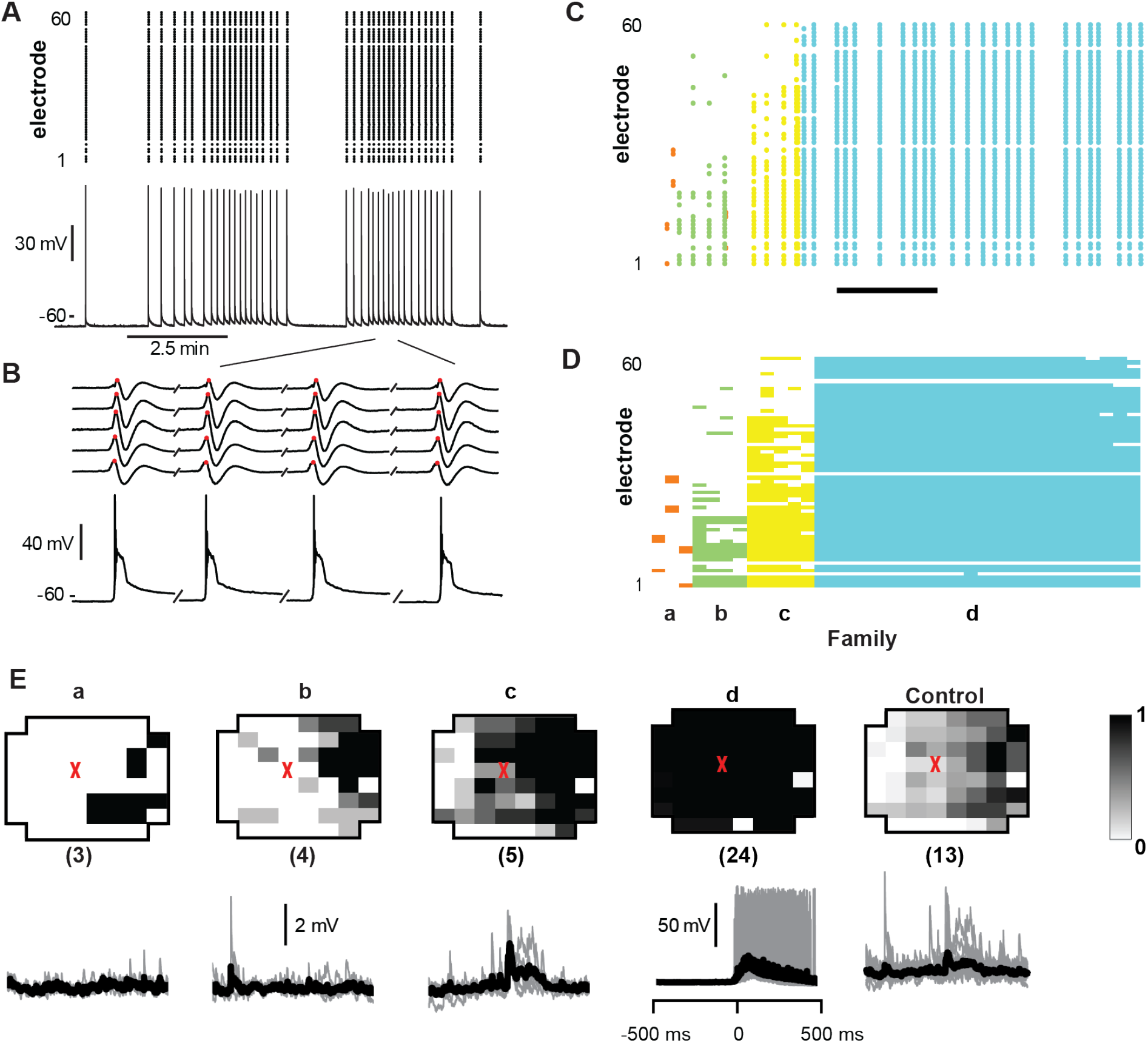
Balanced inhibition is required for selective input. **A.** Sorted population event pattern raster for an example experiment in which PTX (50 μM) was bath applied to block fast GABAA receptor mediated inhibition. Note that population events were stereotyped, most belonging to family *d*. **B.** Average patterns for each family. Red triangle marks soma location of patched neuron. The numbers of trigger events are shown in parentheses. **C.** Family-triggered average membrane potential waveforms (black). Pre-averaged individual traces shown in gray. Vertical scale bars – 2 mV (families *a*, *b*, and *c*), 50 mV (family *d*). Most population events belong to family *d*, which spans most recording sites and always caused the patched cell to fire, i.e. the patched neuron was unselectively involved in network dynamics.

## Discussion

We simultaneously recorded single neuron and multi-site LFP activity from the cortex of awake monkeys and rat acute slices. In both preparations, spatiotemporal LFP patterns distributed in sizes according to a power law, the hallmark of neuronal avalanches. The power law quantifies a high incidence of large LFP patterns suggestive of a non-selective relationship between spatially extended LFP population signals and single neuron activity. On the contrary though, we found that diverse ensembles of extracellular units were selectively and reliably activated with particular repeating avalanche patterns during ongoing activity in premotor cortex of awake monkeys. We confirmed this selectivity in acute slices of rat cortex under two different activity levels, demonstrating reliable input to layer 2/3 pyramidal neurons during select and repeated LFP avalanche patterns. We demonstrated that this selectivity breaks down during disinhibition and is not predicted by the spatially wide-spread correlation found with traditional spike-triggered or LFP-triggered average relationships. The selective participation of single neuron activity with repeated avalanche patterns supports the view that neuronal avalanches are composed of highly diverse, yet selective neuronal ensembles.

Our work is related to previous studies in which multi-site recordings of LFP activity was compared to the activity of single-units (Destexhe et al., 1999;Rasch et al., 2008;Katzner et al., 2009;Nauhaus et al., 2009;Petermann et al., 2009;Kelly et al., 2010). Nauhaus et al. (2009) reported that the spiking activity of neurons generates negative LFP deflections near the neuron and decays with distance from the neuron. This conclusion was based on the spike-triggered average LFP recorded from anesthetized cats and monkeys. Our unit-triggered averages of LFP (Figure 1D) confirm these findings in awake monkeys. However, when lower frequency signals are not filtered out, i.e. 1 – 100 Hz is considered rather than 3-100 Hz as Nauhaus et al. (2009) did, the decay of the spike-triggered average LFP peak with distance is less prominent (Figure 1C). The converse relationship, i.e. LFP-triggered average spike histograms, revealed spatially widespread spiking during negative LFP deflections (Figure 1E). This observation is also consistent with previous observations of nLFP-triggered spike histograms (Destexhe et al., 1999;Petermann et al., 2009). Katzner et al. (2009) found that LFP signals originate from neurons within a 250 μm radius of the recording site. They reached this conclusion by comparing orientation tuning of units and LFP signals in visual cortex of anesthetized cats. Our results do not contradict this study, but we emphasize that neuronal avalanche dynamics is sensitive to anesthetics (Scott et al., 2014;Bellay et al., 2015) limiting extrapolations of previous findings to the current study in awake nonhuman primates. Indeed, previous studies have shown that LFP signals can be highly correlated over many millimeters of cortex (Destexhe et al., 1999;Leopold et al., 2003;Nauhaus et al., 2009). When considered as a whole, our study demonstrates that large average population events involve selective ensembles of units distributed all across the 4 mm × 4 mm sized recording region. Some previous studies investigated the spike-LFP relationship by using spike trains to predict the LFP traces (Rasch et al., 2008) or vice versa (Kelly et al., 2010). Our work suggests that the success of such predictions would be substantially improved if algorithms take into account unit activity far from the LFP recording site as well the multi-site spatial pattern of the LFP.

Our analysis of population events is also related to previous studies using voltage sensitive dye imaging (Tsodyks et al., 1999;Kenet et al., 2003;Han et al., 2008), which provides a spatially extended view of population activity similar to multi-site LFP recordings. As in our study, (Kenet et al., 2003) and (Han et al., 2008) found that population activity patterns repeat during ongoing cortical activity. Similar to our finding, (Tsodyks et al., 1999) showed that a single neuron may fire selectively during certain ongoing ‘preferred cortical states’, which were defined by the spike-triggered average population pattern. However, our results indicate that single neurons are often selective for multiple different population events, not just one ‘preferred cortical state’. Moreover, (Tsodyks et al., 1999) restricted their attention to population events which resemble those caused by sensory stimuli, which, unlike our study, excludes the possibility that a neuron might be selective for an internal cognitive process unrelated to sensory stimulation.

A common view of LFP signals is that their physiological origins are too poorly understood to provide concrete information about cortical dynamics. Our work suggests that this view is due for an update. We show that traditional spike- and LFP-triggered average relationships are much weaker than the fluctuating moment-to-moment spike-LFP relationships. Individual units are not well represented by the ‘average unit’ and individual LFP population events are not well represented by the ‘average event’. When these effects are accounted for, we show that diverse and reliable spiking ensembles underlie the cortical LFP avalanche.

Our treatment of LFP population events was motivated by our studies of ‘neuronal avalanches’ identified in the LFP *in vitro* (Beggs and Plenz, 2003;2004;Stewart and Plenz, 2006;2007;Shew et al., 2009;Shew et al., 2011;Yang et al., 2012), *ex vivo* turtle cortex (Shew et al., 2015) and *in vivo* in the rat (Gireesh and Plenz, 2008) and nonhuman primate (Petermann et al., 2009;Yu et al., 2017;Miller et al., 2019). Our observations that spatial patterns of LFP repeat during ongoing activity was shown previously for neuronal avalanches, but only *in vitro* (Beggs and Plenz, 2004;Stewart and Plenz, 2006). Given the ambiguity about the origin of the LFP within the various experimental settings, it was not previously understood how single neurons were involved with avalanche dynamics identified in the LFP. Our work here demonstrates that neuronal avalanches are underpinned by selective, reliable spiking ensembles of neurons. This selectivity thus allows neuronal avalanches to be proposed as a spatiotemporal organization of Hebbian cell assemblies (Hebb, 1949) lending strong experimental support to a large body of simulations on Hebbian plasticity, neuronal avalanches and criticality (de Arcangelis et al., 2006;de Arcangelis and Herrmann, 2010;Rybarsch and Bornholdt, 2014;Stepp et al., 2015;Hernandez-Urbina and Herrmann, 2017;Michiels van Kessenich et al., 2018;Skilling et al., 2019). By extension, the temporal organization of avalanches (Lombardi et al., 2014;Lombardi et al., 2016) or avalanches within avalanches (Petermann et al., 2009) and corresponding firing patterns of spike avalanches (Ribeiro et al., 2016) might provide a template for Hebb’s ‘phase sequences’.

## Ethics Statement

All experiments were carried out according to the guidelines of the National Institutes of Health (NIH) and the Animal Care and Use Committee of the National Institute of Mental Health (NIMH) and approved animal study protocols LSN-01 and LSN-11.

## Author Contributions

DP, SY, WS and TB designed the research. SY, TB, WS and JWP performed experiments. SY, TB, WS and DP analyzed the data. TB, WS, SY, and DP discussed the results and wrote the manuscript.

## Conflicts of Interest Statement

The authors declare that the research was conducted in the absence of any commercial or financial relationships that could be construed as a potential conflict of interests.

## ACKNOWLEDGEMENTS

The authors thank members of the Plenz lab for discussions and Drs. Richard Saunders (NIMH) and Andy Mitz (NIMH) for help with the nonhuman primate physiology and surgery. This research was supported by the Division of the Intramural Research Program (DIRP) of the National Institute of Mental Health (NIMH) ZIA MH00297.

## supplemental material

No supplementary material.

## Notes

### Competing Interest Statement

The authors have declared no competing interest.

